# Resolving the unsolved: Comprehensive assessment of tandem repeats at scale

**DOI:** 10.1101/2023.05.12.540470

**Authors:** Egor Dolzhenko, Adam English, Harriet Dashnow, Guilherme De Sena Brandine, Tom Mokveld, William J. Rowell, Caitlin Karniski, Zev Kronenberg, Matt C. Danzi, Warren Cheung, Chengpeng Bi, Emily Farrow, Aaron Wenger, Verónica Martínez-Cerdeño, Trevor D Bartley, Peng Jin, David Nelson, Stephan Zuchner, Tomi Pastinen, Aaron R. Quinlan, Fritz J. Sedlazeck, Michael A Eberle

## Abstract

Tandem repeat (TR) variation is associated with gene expression changes and over 50 rare monogenic diseases. Recent advances in sequencing have enabled accurate, long reads that can characterize the full-length sequence and methylation profile of TRs. However, despite these advances in sequencing technology, computational methods to fully profile tandem repeats across the genome do not exist. To address this gap, we introduce tools for tandem repeat genotyping (TRGT), visualization and an accompanying TR database. TRGT accurately resolves the length and sequence composition of TR regions in the human genome. Assessing 937,122 TRs, TRGT showed a Mendelian concordance of 99.56%, allowing a single repeat unit difference. In six samples with known repeat expansions, TRGT detected all repeat expansions while also identifying methylation signals, mosaicism, and providing finer resolution of repeat length. Additionally, we release a database with allele sequences and methylation levels for 937,122 TRs across 100 genomes.

## Introduction

Tandem repeats are regions of the genome consisting of exact or near-exact repetitions of DNA sequence motifs. Many subtypes of tandem repeats have been defined including homopolymers (1bp motifs), short tandem repeats (STRs; 2-6bp motifs), and variable number tandem repeats (VNTRs; >6bp motifs). Tandem repeats contribute a substantial fraction of genetic variation in a typical human genome and are estimated to account for over 70% of structural variants longer than 50 bp (English et al, In Preparation).TR expansions have been linked to over 50 monogenic disorders such as Huntington’s disease ^1^, ALS ^2^ and Fragile X syndrome ^3^. Lengths of many TRs are correlated with gene expression ^4^ and, recently, *de novo* TR expansions have been associated with cancer ^5, 6^ and some neurodevelopmental and psychiatric disorders ^7, 8^. Further, somatic mosaicism of TRs associated with rare disease can affect the age of onset, severity, and progression of disease ^9–11^. Despite this correlation between TR length and phenotype, TRs have been understudied due to the difficulty in developing accurate, high-throughput, genome-wide assays^12^. Additionally, while many bi-allelic variants can be studied indirectly through linkage disequilibrium with SNPs, TRs are likely to be missed in these studies because their hypervariability will tend to eliminate any correlation with surrounding variants^13^. Thus, it is essential to include TRs and their variation in regular genomic studies.

Resolving variation in TR regions is a complex task. A variety of assays have been designed to profile different features of the repeat sequence, including length, specific sequence interruptions, methylation, and mosaicism. Southern blot and PCR-based assays enable a lower-throughput profiling of repeat lengths at a limited number of loci^14, 15^ and detection of repeat interruptions^16^. Recently, informatics methods have been developed to resolve some TRs in short-read sequencing data ^17–24^. These methods make it possible to study repeats at the genome-wide scale, however they are less accurate when the repeat is larger than the sequencing reads (typically 150 bp for short reads). Many known repeats are only pathogenic when their size reaches several hundreds of base pairs ^25^, meaning that short-read sequencing often cannot determine a pathogenic repeat’s exact length and sequence composition. For example, it is not possible to use short-read sequences to reliably distinguish between premutations (165-600 bp) and full expansions (>600 bp) of the *FMR1* repeat ^26^. In such cases, secondary orthogonal testing such as repeat primed PCR or Southern blot is required to determine the length and thus pathogenicity of the repeat. This is a significant limitation when assessing an individual’s genome for pre or full mutation risk alleles.

Because of the length limitations and high structural complexity of many TR regions, many short read STR or TR callers such as GangSTR ^21^ and ExpansionHunter ^19, 22^ focus on tandem repeats that consist of nearly perfect stretches of motif copies. In contrast, long-read sequencing is particularly well suited for comprehensive repeat analysis because it can capture the entirety of the repeat sequence. However, computational methods for analysis of tandem repeats in long reads must also cope with error patterns of the long-read sequencing technologies and the high structural complexity of repeat regions. Recently, a few long-read methods for tandem repeat analysis have been introduced ^27–29^. However, these tools only focus on a few loci or structurally simple repeats, avoiding complex and important TR regions of the human genome. Thus, there is a need for general-purpose methods for tandem repeat analysis capable of profiling both simple STRs and more complex VNTRs. In addition to the basic repeat-length genotyping, a comprehensive analysis of TRs requires tools that can characterize repeat allele sequences, as well as profile and visualize mosaicism. Mosaicism is an inherent feature of cancer-associated genome instability and certain pathogenic repeat expansions. These capabilities are necessary to fully explore the mechanisms and impacts of tandem repeat mutations on disease phenotype.

The high accuracy of PacBio HiFi long-read sequencing now makes it possible to comprehensively characterize both germline and somatic variation of tandem repeats across the genome^30, 31^. Furthermore, the technology enables CpG methylation profiling of TR regions, providing the potential to simultaneously assess genetic and epigenetic mutations of TR regions, reveal hidden patterns and novel biology. In particular, the association between repeat length and methylation status can be leveraged to detect highly methylated pathogenic expansions. For example, individuals with reduced methylation of the *FMR1* repeat have been observed to have a reduced Fragile X syndrome phenotype^32^. Currently, we need a method that combines these signals to fully leverage the potential of PacBio HiFi sequencing in revealing new key insights into TR regions at scale.

Here we describe the Tandem Repeat Genotyping Tool (TRGT), a novel method for repeat analysis of long reads, as well as a companion method for Tandem Repeat Visualization (TRVZ). TRGT makes it possible to analyze structurally complex tandem repeats that cannot be accurately represented by other available methods. Additionally, TRVZ affords a visual inspection of repeat alleles. Visualization is especially important when assessing clinically important repeats and is recommended by the Association for Medical Pathology and the College of American Pathologists ^33^. TRGT reports haplotype-resolved germline variation together with methylation status across simple and complex TRs, and can detect mosaic mutations.

## Results

### Accurate Tandem Repeat variant calling using PacBio HiFi DNA sequencing data

TRGT is designed for analysis of repeat alleles in HiFi sequencing data across a user-provided list of repeat regions. The input for TRGT consists of a BAM file with aligned HiFi reads and a list of repeat definitions (**Figure 1A**). Briefly, TRGT works by locating the repeat flanks in each read overlapping the repeat (**Figure 1B**), clustering the reads to determine the consensus sequence for each repeat allele (**Figure 1B**) and then using the repeat structure defined for each locus (**Figure 1C**) to locate the boundaries of motif copies within each allele (**Figure 1D**). While the structure of simple repeats is defined by specifying the repeating motif, more complex repeats are defined by hidden Markov models (HMMs) (**Figure 1C**), following earlier work that demonstrated suitability of HMMs for modeling TRs ^34^. The output of TRGT consists of a VCF file ^35^ with annotated repeat allele sequences (**Figure 1E**) and their methylation levels. TRVZ is a companion tool to visualize the reads aligned to the repeat alleles (**Figure 1F**) and can be used to determine the accuracy of the genotype results returned by TRGT.

**Figure 1:**
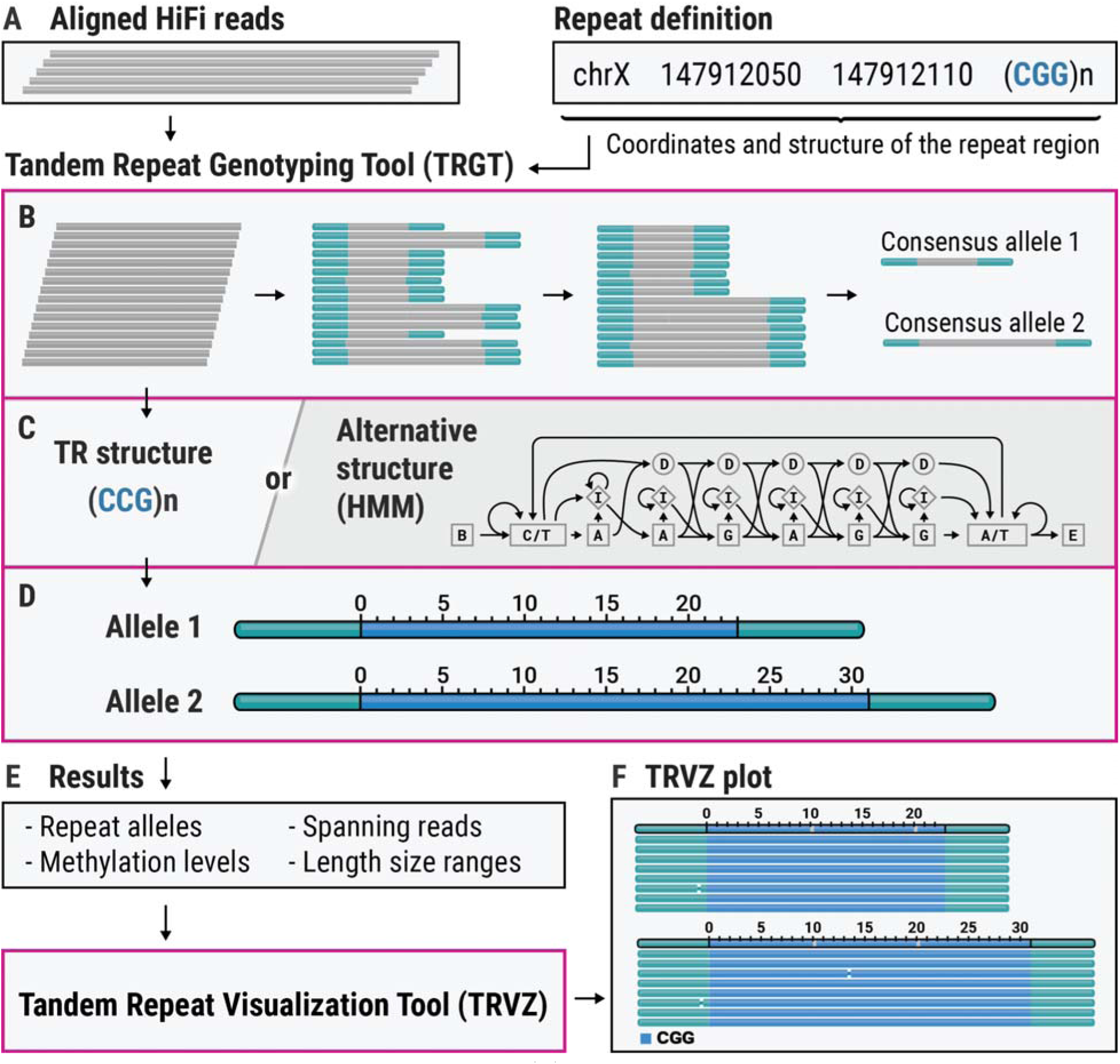
An overview of TRGT and TRVZ. (A) Input to TRGT consists of HiFi reads and a list of repeat definitions. (B) TRGT determines consensus repeat alleles. (C-D) TRGT uses the pre-specified structure of the tandem repeat region to locate individual motif copies in each repeat allele. More complex repeat regions are specified with hidden Markov models. (E) Overview of key fields in TRGT’s output. (F) TRVZ generates plots that display repeat alleles and reads aligning to them, with optional methylation.

To assess the accuracy of TRGT genotype calls, we used TRGT to genotype 937,122 repeat regions spanning 122 Mbps of the reference genome ^36^ in 36x whole genome HiFi sequencing of HG002 from the Genome in a Bottle. We then compared the resulting repeat alleles to a recent state of the art assembly of the same sample from the “Telomere-to-Telomere” (T2T) Consortium ^37–39^. Compared to this assembly, 97.70% of the alleles either agreed exactly or had at most a single base pair difference (**Figure 2C**). To further assess the accuracy of the genotypes, we calculated the Mendelian consistency of repeat lengths across the family trio consisting of HG002, HG003, and HG004 samples (**Figure 2A**). Overall, TRGT showed a Mendelian consistency rate of 88.83% across all repeats and most of the errors (96.06%) corresponded to cases where the number of repeats differed by one between a parent and child (**Figure 2B**). Ignoring such “off-by-one” calls results in a Mendelian consistency rate of 99.56%. As expected, homopolymers and dinucleotide repeats were more error prone (81.85% exact and 99.39% off-by-one consistency) compared to repeats with longer motifs (98.02% exact and 99.78% off-by-one consistency). TRGT completed the analysis across these 30-fold sequencing coverage datasets in 35 minutes using 32 CPU cores.

**Figure 2:**
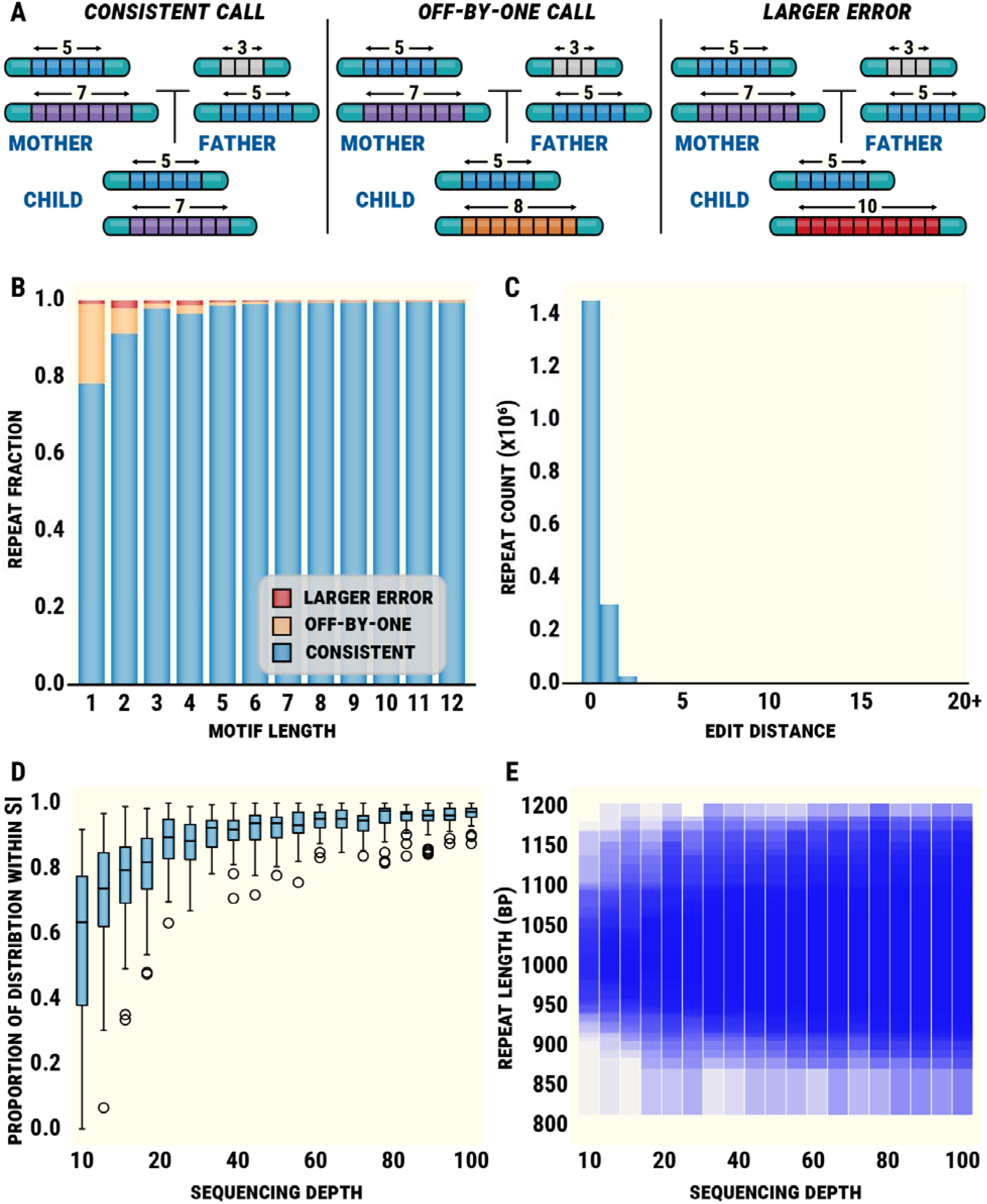
TRGT benchmarks. (A) Examples of a consistent genotype, an off-by-one error, and a larger error. (B) A histogram stratifying the distribution of Mendelian errors by motif length. (C) Edit distances between repeat alleles estimated by TRGT and an HG002 genome assembly. (D) The proportion of the expanded FMR1 repeat distribution captured by TRGT’s size intervals from subsampled 500X NoAmp targeted sequence, and (E) the density of TRGT’s size intervals.

In addition to TRGT, we evaluated two other methods used for profiling tandem repeats in long-read sequencing data, tandem-genotypes ^28^ and Straglr ^29^, and also one method designed for short reads, GangSTR^21^. The same repeat catalog was used for all methods. Because these tools are only designed to estimate repeat lengths and not repeat sequence or mosaicism, we assessed them by measuring length-based Mendelian consistency. TRGT genotyped 99.17% of repeats in all family members while tandem-genotypes, Straglr, and GangSTR genotyped 98.34%, 59.15%, and 95.07% of repeats, respectively. Mendelian consistency allowing off-by-one calls was 98.54%, 83.13%, and 76.60% respectively for Tandem-genotypes, Straglr, and GangSTR compared to 99.56% for TRGT. Thus, TRGT represents a substantial improvement in accurately measuring tandem repeat length. Furthermore, TRGT can assess sequence context, measure repeat methylation, mosaic changes, and facilitate repeat visualization via TRVZ.

Next, we assessed TRGT’s ability to detect mosaic expansions where, instead of a single expanded allele, we observe reads supporting a distribution of allele sizes falling within a certain size range. For this analysis, we focused on the *FMR1* repeat region in the NA07537 sample, which was sequenced to nearly 500x HiFi read depth using the NoAmp targeted assay ^40^. TRGT estimated that the length of the mosaic expansion ranges from 813 to 1204 bp, which was concordant with the previous studies of this sample^41^. To assess TRGT’s ability to accurately determine mosaicism at lower sequencing depths, we subsampled these data to depths ranging from 10x to 100x (100 replicates were performed at each depth). We then measured the proportion of the expanded alleles observed in the original sample captured by the corresponding TRGT’s allele size interval. On average, over 75% of the expanded alleles were captured at 15x sequencing depth or higher and, as expected, the confidence intervals were centered at 1000 bp – the point estimate of the expansion size (**Figure 2D & E**).

### Population analysis of tandem repeats

To study the genome and population-wide variability of the 937,122 TRs, we built the TRGTdb database (Methods) from a collection of 104 HiFi samples from Genome in a Bottle (GIAB) and the Human Pangenome Reference Consortium (HPRC) (**Table S1**). We measured the length polymorphism of a repeat by computing the number of its alleles of distinct lengths per 100 samples (length polymorphism score). To reduce conflation between true alleles and technical artifacts (e.g., one-off errors in homopolymer regions, **Figure 2B**), we only used alleles appearing at frequency above 1% (i.e., observed at least three times). We observed that 13.64% of alleles showed no evidence of recurrent mutations (3.64% were monoallelic and 10% were biallelic), while 86.36% were multi-allelic. Of the multi-allelic loci, 61.85% had 3-5 alleles, 27.71% had 6-10 alleles and 10.45% had more than 10 alleles (**Figure 3A**).

**Figure 3:**
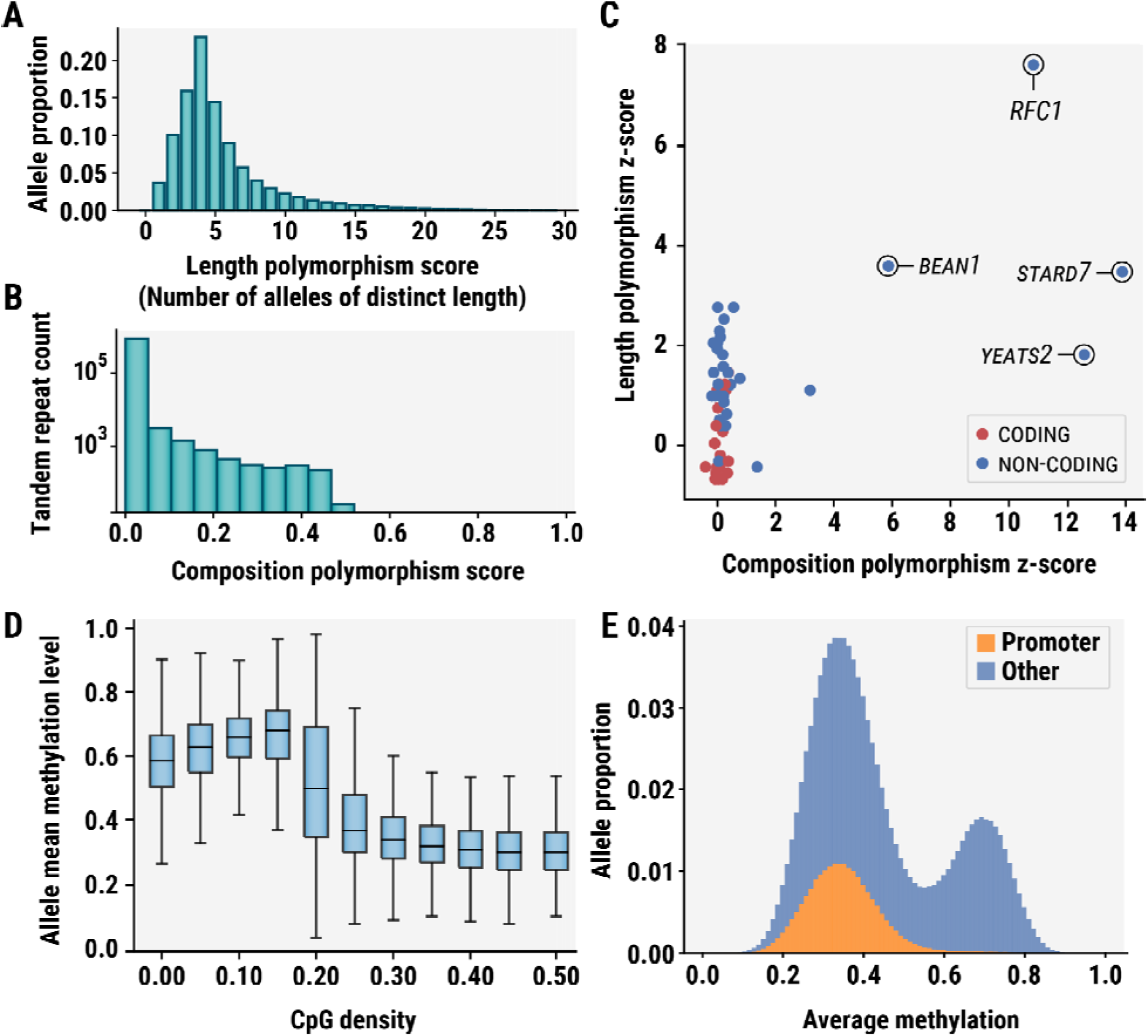
Genetic and epigenetic variation in TR regions across 104 HPRC samples. (A) Distribution of length polymorphism scores defined as the number of alleles of distinct length per100 samples. (B) Distribution of allele composition polymorphism scores. (C) Length and composition z-scores for known pathogenic repeats. (D) Distribution of allele mean methylation levels stratified by CpG density. (E) Mean methylation levels of TRs overlapping CpG islands.

We investigated variations in the composition of the repeat sequences. For this, we compared the composition of two repeat alleles by calculating one minus the Jaccard Index between the corresponding sets of high-frequency k-mers that we call the composition difference score (Methods). For example, the composition difference score of repeat alleles CAG * 10 and CAG * 100 is 0.0 because of their identical composition despite the significant length difference. In contrast, although alleles CAG * 10 and CAA * 10 have the same length, their composition difference score is 1.0. To measure the degree of sequence composition polymorphism of a repeat, we calculate the mean of composition difference score for all pairs of repeat alleles. We refer to this value as the composition polymorphism score (CPS) of the repeat. The CPS was below 0.01 for over 98% of repeats, and only 0.31% of repeats had composition polymorphism scores above 0.2 (**Figure 3B**). This distribution indicates that tandem repeats tend to have very homogenous sequence composition.

Given this collection of samples, we next characterized the variation of known pathogenic repeats. To compare the length and composition polymorphism of known pathogenic repeats in HPRC samples relative to our genome-wide repeat catalog, we calculated z-scores for the length and composition polymorphism scores for 56 known pathogenic repeats relative to the corresponding genome-wide distributions (**Figure 3C**). Consistent with our expectations, coding pathogenic repeats exhibited little polymorphism (**Figure 3C**). In contrast, non-coding repeats tended to have higher length polymorphism compared to other repeats across the genome. Furthermore, *STARD7*, *YEATS2*, *RFC1*, and *BEAN1* were the only pathogenic loci to exhibit substantial composition polymorphism. This is consistent with the fact that pathogenic expansions of these repeats correspond to changes in the motif sequence. Our analysis suggests that studies focused on identifying novel pathogenic expansions may prioritize non-coding repeats with polymorphic length or composition in addition to coding repeats.

We also profiled CpG methylation by using TRGT to estimate the mean methylation level of each repeat allele. The resulting distribution of methylation levels was consistent with the expected human genome methylation profile: CpG denser regions had markedly lower methylation compared to the CpG sparser regions (**Figure 3D**).

We next focused our analysis on TR loci that overlap CpG islands and annotated each by their intersection with promoters ^42, 43^. In total, 9,821 TR loci overlap CpG islands and 2,671 overlap promoters The average methylation levels of 1,425,694 TR alleles overlapping CpG islands have a bimodal distribution (**Figure 3E**). The lower peak of the distribution can be partially explained by CpG island TR alleles overlapping promoters. These findings confirm previous observations on the relationship between CpG islands, promoters, and methylation ^44, 45^. When considering all TR alleles which fall within the top third of the average methylation range (corresponding to methylation levels between 0.68 and 1.0), we found that only 0.5% overlap with CpG islands. Moreover, we identified 2,315 alleles originating from 552 loci that overlap promoters and exhibit an average methylation level greater than 0.68. Among these 552 loci, 317 had more than two observed alleles with higher average methylation.

We next analyzed the genomes of six individuals sequenced at Children’s Mercy Kansas City who were previously identified to carry repeat expansions at known pathogenic loci in one of four genes: TRGT correctly identified the pathogenic expansions in each sample, calling an *FMR1* 350 bp premutation and 1 Kbp full-expansion (117 and 300 CGG motifs, respectively), a *DMPK* expansion spanning over 5 Kbp, two *STARD7* expansions each spanning over 1 Kbp, and an *ATXN10* expansions >4 Kbp (**Table S2, Figure S3**). Compared to the previously applied testing that only sized the repeat to broad ranges, TRGT identified the size of the repeats to nearly bp resolution.

### Detailed characterization of *RFC1* repeat region

A repeat region within the *RFC1* gene located at chr4:39348424-39348479 (hg38) has been recently associated with cerebellar ataxia, neuropathy, and vestibular areflexia syndrome (CANVAS) ^46, 47^. Unlike most other pathogenic repeats, *RFC1* repeat alleles are known to have heterogeneous sequence composition consisting of stretches of AAAAG, AAAGG, and other motifs. CANVAS itself has been linked to biallelic expansions consisting of either AAGGG or ACAGG motifs. Within TRGT, this repeat is described by a hidden Markov model (HMM) whose topology is defined by the constituent motif sequences (**Figure 4A**). The HMM makes it possible to segment the sequence of each allele into a set of regions spanned by each motif. For example, the short allele of *RFC1* repeat in the HG00733 sample consists of stretches of AAAGG, AACGG, GGGAC, and AAGGG motifs, while the long allele consists of a 600 bp stretch of AAAAG motif (**Figure 4B**). To investigate the population-level structure of *RFC1*, we used TRGT to analyze this repeat in the 104 HPRC samples. We first summarized the composition of each allele by computing the fraction of its sequence spanned by each constituent motif (**Figure 4C**). This allowed us to group the alleles into six composition clusters (**Figure 4C,D**). The alleles in each cluster are characterized by the presence of a relatively long stretch of one of the five motifs (AAAAG, AAGAG, AAAGGG, AAGGG, AACGG, and GGGAC). The largest cluster consisted of alleles composed of the AAAAG motif. The alleles within this cluster could be further subdivided into two groups: short alleles spanning less than 200 bp and longer expansions spanning over 300 bp (**Figure 4E**). Another cluster consisted of three alleles containing long stretches of the pathogenic AAGGG motif (**Figure 4C,D**), which is consistent with the high carrier frequency of this expansion ^46–49^.

**Figure 4.**
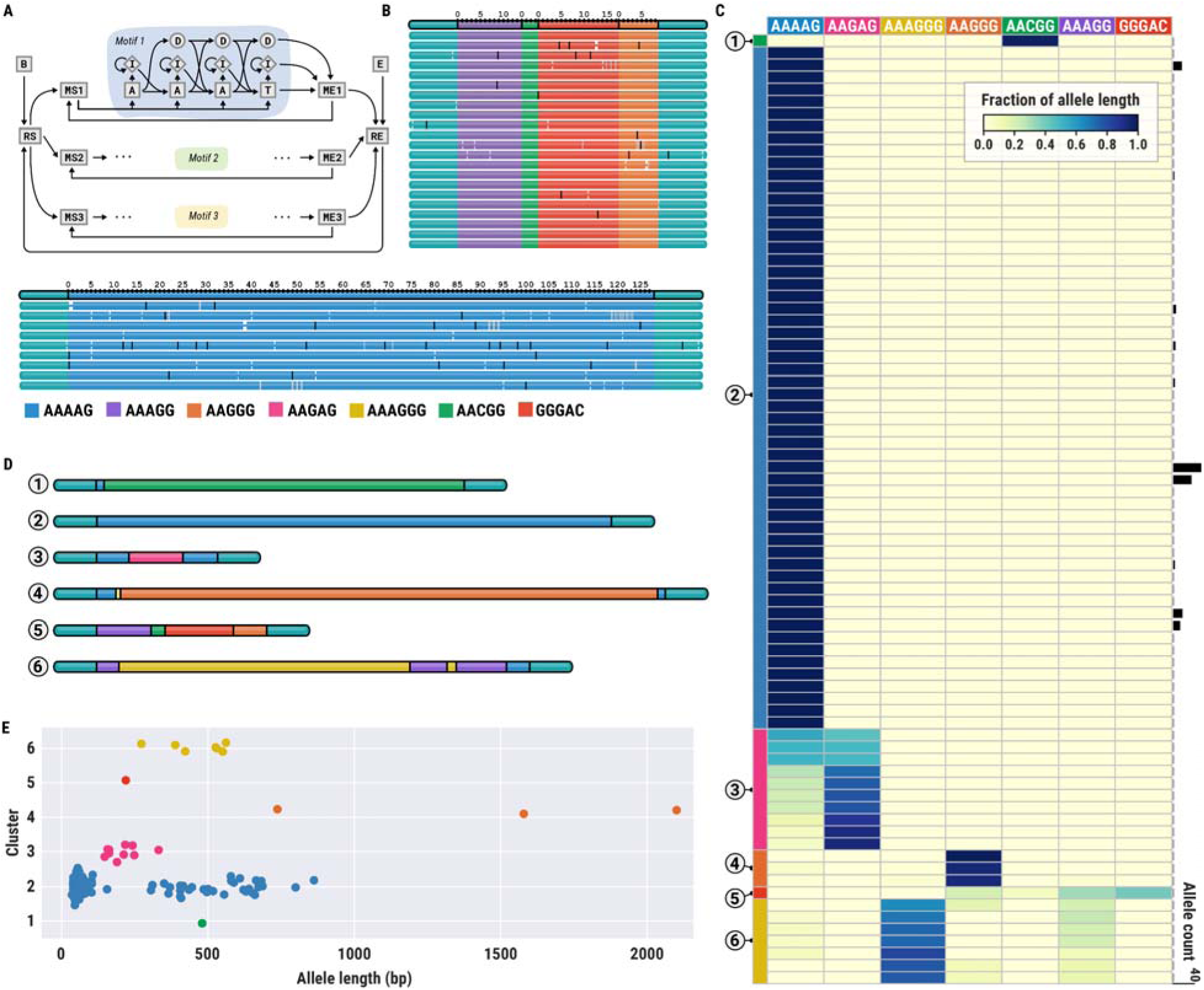
Genetic variation of RFC1 repeat alleles. (A) A hidden Markov model representing the population structure of the RFC1 TR. (B) A TRVZ plot depicting both alleles of the RFC1 repeat in the HG00733 sample. (C) A heatmap depicting the span of each motif (columns) on each allele (rows); each cluster of alleles is associated with the color of its dominant motif. (D) An example allele from each cluster. (E) Lengths of alleles belonging to each cluster.

### Analysis of complex FMR1 extension for non, partial, and full mutation carriers

We analyzed the CGG repeat in the promoter region of the *FMR1* gene,associated with Fragile X syndrome^50^. *FMR1* alleles containing between 55 and 200 CGGs are called premutations and have been linked with fragile XLJassociated ataxia syndrome and fragile XLJassociated primary ovarian insufficiency ^50^. Alleles with 200 or more CGGs are called full expansions and cause Fragile X syndrome. Full expansions have been associated with heavy CpG methylation as well as mosaicism, meaning that the exact length of the expanded repeat can vary across the cells ^50^. In addition to the overall length, AGG interruptions within the repeat sequence have been associated with increased stability of the repeat and reduced the risk of a parent with a premutation passing a full expansion to their child ^51^.

The overall distribution of *FMR1* allele lengths (**Figure 5A**) was similar to that reported previously, with a mean size of ∼30 repeats ^52^. Our analysis also identified two *FMR1* alleles of premutation length in HG04184 (58 motifs) and HG00438 (60 motifs) samples (**Figures 5B** and **5C**, respectively).

**Figure 5:**
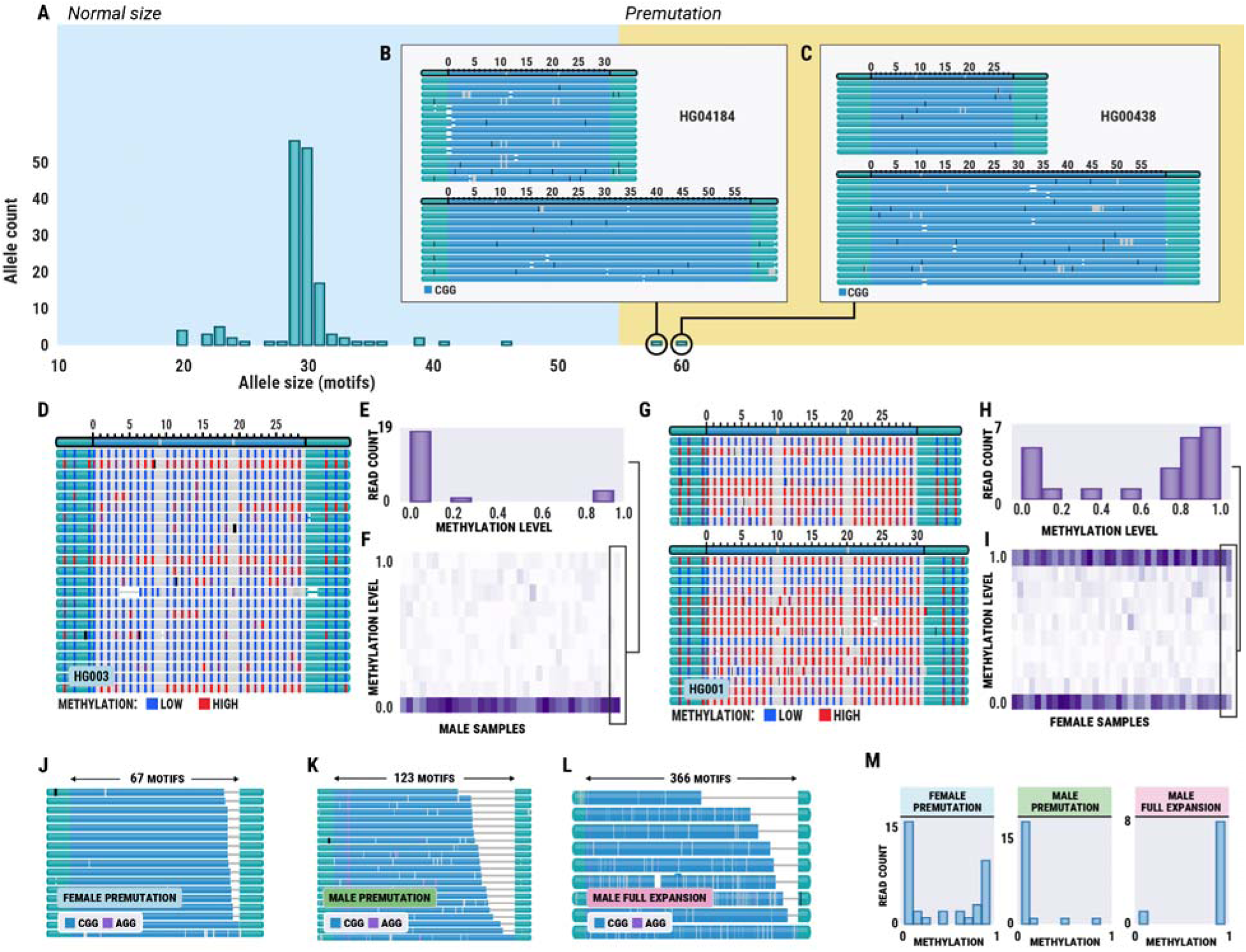
Genetic and epigenetic variation of FMR1 repeat. (A) Distribution of FMR1 allele sizes in 100 HPRC samples. (B,C) TRVZ plots of FMR1 repeat in the HG04184 and HG00438 samples, respectively, showing premutation alleles. (D) TRVZ plot of FMR1 repeat in the HG003 male sample displaying CpG methylation. (E) Distribution of median methylation levels for HG003 reads spanning FMR1 repeat. (F) Distributions of median methylation levels for FMR1 reads across all male samples. (G) TRVZ plot of FMR1 repeat in HG001 female sample displaying CpG methylation. (H) Distribution of median methylation levels for HG001 reads spanning FMR1 repeat. (I) Distributions of median methylation levels for FMR1 reads across all female samples. (J) Premutation repeat allele from a prefrontal cortex sample from a female donor (short allele not shown). (K) Premutation repeat allele from a prefrontal cortex sample from a male donor. (L) Fully expanded repeat allele from a prefrontal cortex sample from a male donor. (M) Methylation profile of prefrontal cortex samples.

As most of the X chromosome in males is transcriptionally active, we expected *FMR1* methylation to be low. Indeed, the vast majority of reads spanning this repeat in the HG003 male sample were almost completely devoid of methylation (**Figure 5D, E**). We observed the same low methylation pattern in all other male samples that we analyzed (**Figure 5F**). Next, we analyzed the HG001 (NA12878) sample derived from a healthy female donor. Both alleles of this repeat spanned significantly fewer than 55 CGG copies and two AGG interruptions, conclusively indicating that this individual is not a carrier. We observed a bimodal methylation pattern of each allele consistent with chromosome X inactivation (**Figure 5G**). Summarizing the distribution of median methylation levels for each read in this (**Figure 5H**) and all other female samples (**Figure 5I**) confirms the bimodal nature of *FMR1* methylation in females.

Finally, we analyzed the FMR1 repeat in three brain samples previously determined to have the pre-mutation (two samples) or full mutation (one sample). TRGT estimated the samples with expected premutations to have an *FMR1* repeat spanning 67 and 123 copies of the repeat motif (**Figure 5J, K**), consistent with the established range for premutations. Interestingly, one of the premutation samples did not contain the stabilizing AGG interruptions (**Figure 5J**), signaling an increased risk of transmitting a full expansion to children. The sample with expected full mutation was estimated to contain 325 motif copies and exhibited a strong degree of mosaicism with repeat lengths ranging from 200 to 366 (**Figure 5L**). Importantly, all samples showed the expected methylation patterns (**Figure 5M**). The female pre-mutation sample (Sample 5) exhibited bimodal methylation, whereas the male pre-mutation sample (Sample 2) was lowly methylated. In contrast, the male sample with the full expansion was highly methylated, characteristic of the Fragile X syndrome. These results demonstrate the utility of TRGT and TRVZ to accurately identify and visualize complex TRs alongside patterns of mosaicism and methylation across different samples and tissue sources.

## Discussion

We described a new software tool, TRGT, to quantify tandem repeats from HiFi sequencing data and demonstrated that it can accurately characterize both known pathogenic repeats and a genome wide catalog of almost one million TRs. Compared to the other long-read TR methods, TRGT achieves significantly higher Mendelian consistency of 98.02. Mendelian consistency rose to 99.78 if off-by-one errors are excluded. In addition to accurately genotyping TRs, we have included two companion methods to increase the utility of TRGT. 1) TRVZ allows users to visualize the read level evidence supporting the genotype calls made by TRGT, and 2) TRGTdb builds a database of TRs that can be used to annotate sample specific variant calls relative to a population. These companion methods will help researchers and clinical labs annotate and visually inspect the genotypes made by TRGT.

Compared to current methods for testing known pathogenic repeats, TRGT combined with HiFi reads can, as a single test, deliver many features that match or even surpass the performance of current sequencing techniques in general wet-lab based testing protocols. For example, TRGT provides an exact motif count of a repeat. This is especially critical for assessing individuals carrying pathogenic repeats. For affected individuals, TRGT provided an exact count of the numbers of repeats and indicated the range of mosaicism. In contrast, other established sequencing or wet lab methods (e.g. repeat-primed PCR and Southern blot) merely provide size ranges or an average repeat length. Additionally, TRGT quantifies both the size and the motif sequence of repeats as is which is critical to interpreting loci such as *RFC1* and *SAMD12*^53^. Finally, because TRGT also reports the average methylation from HiFi sequence reads, users can get the repeat length, sequence context and methylation status from a single sequencing experiment. While TRGT can provide all of this information as a single test, it should be noted that because it requires spanning reads, it may fail to call variants in low sequencing depth samples or regions. Efforts are currently underway to improve TRGT’s ability to identify pathogenic repeat expansions with lower sequencing coverage or in cases when reads do not fully span the repeat, as can happen for particularly large expansions.

Most of the known repeats become pathogenic at sizes beyond what can be resolved with short reads alone^54^. For example, pathogenic *FMR1* expansions are >200 repeats (>600 bp) and size estimates for *FMR1* are consistently underestimated even when using state of the art short read repeat callers ^26^. Additionally, TRs with high mutation rates are unlikely to exhibit linkage disequilibrium with surrounding SNPs ^13^. This means that SNP-based studies will be unlikely to detect these TR risk alleles through association. Conversely, a genome wide catalog of TRs genotyped with TRGT, possibly sequenced at lower depths, will greatly improve the power to detect TRs associated with complex traits. Because tandem repeats may be more likely to have epistatic interactions, association studies that include accurate genotyping of all variants including TRs may help explain much of the missing heritability^12^.

Though TRGT includes many features absent from other TR-specific variant callers, there are areas for continued development. While we cataloged almost a million repeats across 100 samples, there is a significant need to extend this database to genotype more TRs and include more samples of diverse ancestry and detection of novel loci. With a more complete and diverse database, we can perform a more systematic analysis of repeat length and sequence context as well as methylation levels. This database can be leveraged to identify whether TRs in a sample are significantly expanded relative to the population in much the same way that frequency databases like gnomAD^55^ are used to annotate SNPs or indels. We are also continuing to extend our repeat catalog to include a more complete representation of all variable repeats including ones that may not show up as repetitive in the current reference genome.

We showed that the tools TRGT, TRVZ, and TRGTdb can highlight many important properties that are observed in known pathogenic TRs, including hyper methylation and variability in the repeat sequence. This demonstrates a significant advance in the tools available for unraveling the often under-explored complexity of tandem repeats. Combined, these tools will enable researchers to gain novel insights on many aspects of evolution, genetic diversity, and the medical implications of TRs.

## Methods

### Tandem Repeat Genotyping Tool (TRGT)

TRGT performs tandem-repeat genotyping using HiFi reads that overlap each repeat. The input to TRGT consists of a BAM file ^56^ with aligned HiFi reads and a file with repeat definitions. The output consists of a VCF file containing full-length repeat alleles and their methylation levels as well as a BAM file with portions of HiFi reads that span each repeat. Analysis of each repeat region proceeds as follows:

1. TRGT locates reads that span a given repeat region and these reads are assigned to each allele by clustering. To cluster the reads, TRGT first calculates the edit distances between all pairs of reads and then performs agglomerative clustering using Ward linkage ^57^. We then filter out any cluster containing fewer than 10% of the total number of spanning reads. For diploid repeats, we assign the two largest clusters to each allele. For haploid repeats we assign the largest cluster to each allele.
2. To determine the consensus sequence of each repeat allele, TRGT selects a read of the median length from the corresponding cluster of reads and uses it as the alignment backbone. All reads in the cluster are aligned against this backbone sequence. The consensus sequence is then determined by scanning the backbone and incorporating bases by performing a majority vote on the alignment operations. For example, if most read alignments contain a sequence insertion at some position of the backbone, then this sequence is incorporated into the consensus.
3. TRGT next annotates occurrences of individual repeat motifs within the sequence of each consensus allele. Different annotation algorithms are used depending on the type of repeat. For example, simple tandem repeats that can be described by repetitions of one or multiple fixed-sized motifs are annotated using a fast algorithm based on finding the longest path in an acyclic graph. More complex repeats are annotated using hidden Markov models. These annotation methods are described below.
4. The methylation level of each repeat allele is set equal to the mean methylation level of all CpGs in the corresponding region in all reads that support the allele.

### Annotation of simple tandem repeats regions

We define **simple TR regions** as regions whose population structure can be described as a series of TRs, possibly separated by interrupting sequences. The structure of such regions is described by an expression (m)^nl^s (m)^n^^2^ ..s(m)^nk^ where m is the motif of ith TR, n is the (allele-specific) motif count of ith TR, and s_i_ is a possibly empty sequence separating TR i and i+ 1. Given a query allele sequence (**Figure S4A**), the segmentation algorithm proceeds as follows. First, we create a graph whose nodes correspond to matches between the query sequence and motifs m_i_ and interrupting sequences s_i_ (**Figure S4B**). Then, we create a directed edge from node m_i_ to node x if x is the next occurrence (in the topological order induced by the query sequence) of m_i_, s_i_, or m_i+1_ (**Figure S4C**). Nodes s_i_ are connected using the same rule. We then determine a path that spans the largest number of bases. This path can be determined by calculating the longest path in a directed acyclic graph, which covers the largest number of bases terminating at each node (**Figure S4D**). This path corresponds to the segmentation of the original query sequence (**Figure S4E,F**).

### Annotation of complex tandem repeat regions

Certain tandem repeat regions cannot be represented by the expressions introduced in the previous section. We call such regions complex following previous work ^34^. We use hidden Markov models to model the structure of these repeats. TRGT can synthesize HMMs that model sequences that correspond to runs of a specified set of motifs. These runs can occur in an arbitrary order. *RFC1* (**Figure 4**) is one example of such repeats. HMMs of this family all have similar topology (**Figure S5**): The customary start and end states (**Figure S5A,B**); a pair of silent states delineating the start and end of each motif run (**Figure S5C,D**); a pair of states delineating the start and end of each repeat motif (**Figure S5E,F**); and finally a block of states representing the motif occurrence sequence consisting of states corresponding to matches / mismatches, insertions, and deletions of motif bases. TRGT can also accommodate HMMs with other topologies. In this case, it requires that the HMM specification includes a list of edges that connect the terminal nodes of each motif as well as the sequence or label of each motif.

### Tandem Repeat Visualization (TRVZ) tool

TRVZ is a companion visualization tool for TRGT allowing users to view selected repeats of interest. The input to TRVZ consists of files generated by TRGT. The output is an image in either svg, pdf, or png file formats. TRVZ generates a read pileup plot corresponding to each repeat allele (**Figure S6**). The top track of each allele plot shows the consensus sequence determined by TRGT (**Figure S6A**). The consensus is annotated according to its alignment to the perfect repeat of the same length. The solid color corresponds to matches, gray blocks to mismatches, horizontal lines to deletions, and vertical lines to insertions in the allele sequence relative to the perfect repeat. For example, two AGG interruptions that are typically present in the sequence of non-expanded *FMR1* repeats will result in two mismatches (**Figure S6**) because this repeat is defined as (CGG)^n^. The tracks below the top track correspond to the alignments of HiFi reads to each repeat allele (**Figure S6B**).

### TRGT database

Variant Call Format (VCF) entries are useful for representing variation but can be difficult to leverage for programmatic queries of the data. To normalize the data contained within VCFs, we consider each VCF entry to contain information that can be split into three tables: Locus, Allele, Sample. The Locus information corresponds to the VCF entry’s CHROM and POS columns which represent a location in the reference. The Allele information corresponds to the VCF entry’s ALT, QUAL, FILTER, and (generally) INFO columns which represent variation observed at a Locus. The Sample information corresponds to the VCF entry’s FORMAT and SAMPLE columns which represent descriptions of Alleles observed in a sample at a Locus. VCF information is extracted and held in-memory as three pandas DataFrames (one for each table) before being saved on-disk using Apache parquet. Apache parquet is an efficiently compressed, column-oriented file format. To store information across multiple runs of TRGT, all Loci and Alleles are consolidated into a single table and stored in their own parquet file. However, each Sample is stored in its own parquet file. These files are organized within a directory representing the database. By storing each table separately, the genotype information can be removed via deletion of Samples’ parquet files. Full de-identification can be achieved with a TRGTdb command for removing allele sequences, randomizing allele numbers, or shuffling genotypes across samples. On average, a single sample from the 104 sample HPRC TRGTdb has an on-disk storage size of 11.4Mb using TRGTdb compared to individual bgzip compressed VCFs requiring 92.3Mb, an 87.6% decrease.

To assist users with creating a TRGTdb, command line tools are distributed as part of the TRGT package. Command line tools for ‘standard’ queries are included, such as allele counts, the number of monozygotic reference sites, and per-locus genotype information. The outputs of these queries can be saved in tab-delimited, comma-separated, parquet, or joblib formats.

Finally, to assist users in creation of custom queries, a TRGTdb python API is also distributed. Full documentation on the TRGTdb tool is available at (https://github.com/ACEnglish/trgt/blob/main/tdb_tutorial.md). Annotation of TR loci within the TRGTdb against UCSC genome tracks was performed using PyRanges (ref https://academic.oup.com/bioinformatics/article/36/3/918/5543103). All analyses performed with TRGTdb can be recreated by following the jupyter notebook tutorials available (link to notebooks).

### Tandem repeat benchmark

To assess TRGT’s sensitivity to expanded pathogenic STR loci, we ran it on WGS of six individuals with orthogonally confirmed clinical assays. These individuals were enrolled in the Genomic Answers for Kids program^58^. Samples were collected and sequenced on PacBio HiFi Sequel II and IIe systems as previously described ^59^. Sex was inferred using Somalier ^60^ then provided to TRGT using the --karyotype flag. TRGT v0.4.0 was run at known pathogenic loci, using pathogenic_repeats.hg38.bed (commit b10e7f5). Expansions identified by TRGT were further visualized using TRVZ v0.4.0 (**Figure S3**). Orthogonal clinical testing was performed by triplet-primed PCR or Southern blot as part of clinical care (**Table S2**). The subsampling analysis was performed by randomly selecting reads from the original BAM file to achieve the desired depth and then applying TRGT/TRVZ to the resulting subsampled BAM file.

We compared TRGT calls to those made from a high quality assembly. We compared TRs to the HG002 diploid genome assembly as follows: (1) we extracted sequences of all repeat alleles from the HG002 VCF file generated by TRGT (2) we added a 250 bp flanking sequence to both sides of each allele (extracted from the HG38 reference genome) and mapped the resulting sequences to the paternal and maternal contigs of HG002 assembly with minimap2, (3) we picked the top scoring assignment of alleles to paternal contigs for each TR. The benchmarks used Straglr v1.4.1 ^29^, GangSTR v2.5.0 ^21^, and tandem-genotypes v1.9.0 ^28^. TRGT was ran with default parameters; tandem-genotypes was ran with parameters “-o2 --min-unit=1”; Straglr was ran with parameters “--min_str_len 1 --max_str_len 1000 --max_num_clusters 2”; GangSTR was ran with default parameters. Mendelian consistency analysis was performed by genotyping the repeats in the HG002, HG003, and HG004 family trio with each method and then comparing the lengths of repeats in the child to those of their parents (**Figure 2A**). Fractional lengths were rounded to the nearest integer.

### TR composition analysis

To study the variation in sequence composition of TR alleles, we first defined the composition difference score (CDS) that compares sequences of two alleles. Then we used CDS scores to define the composition polymorphism score (CPS) that measures the variation in sequence composition of a TR across a given set of samples. The CDS score between alleles a_1_ and a_2_ is defined by:

> *CDS(a_1_, a_2_, k, n) = 1 - Jaccardlndex(S(a_1_, k, n), S(a_2_, k, n)),*

where S(a_i_, k, n) is the set of k-mers of length k present in the allele a_i_ that appear at least n times in at least one repeat allele. The Jaccard index between two sets is defined as the size of the intersection of these sets divided by the size of their union. For our analyses we used k-mers of length 5 that appear 5 times or more times in at least one allele (k = 5 and n= 5). We then defined CPS score for a TR as the mean of CDS scores calculated for all pairs of alleles.

## Data availability

Version 0.7 of the HG002 assembly from the “Telomere-to-Telomere” (T2T) Consortium was downloaded from <u>GitHub</u>. The data created as part of Genomic Answers for Kids is available through <u>NIH/NCBI dbGAP</u>. Human Pangenome Reference Consortium (HPRC) data is available at NCBI SRA under the <u>BioProject IDs PRJNA850430</u>. The short-read data for HG002, HG003, HG004 is available from the <u>1000 Genomes Phase 3 Reanalysis with DRAGEN 3.5 and 3.7</u> within the Registry of Open Data on AWS.

TRGT, TRVZ and TRGTDB binaries and loci definitions: https://github.com/PacificBiosciences/trgt

### Institutional review board approval

The institutional review board of Children’s Mercy Kansas City (Study#11120514) approved this study.

## Acknowledgements

We thank generous donors to Genomic Answers for Kids program at Children’s Mercy Kansas City.

## Supplementary information

**Figure S1:**
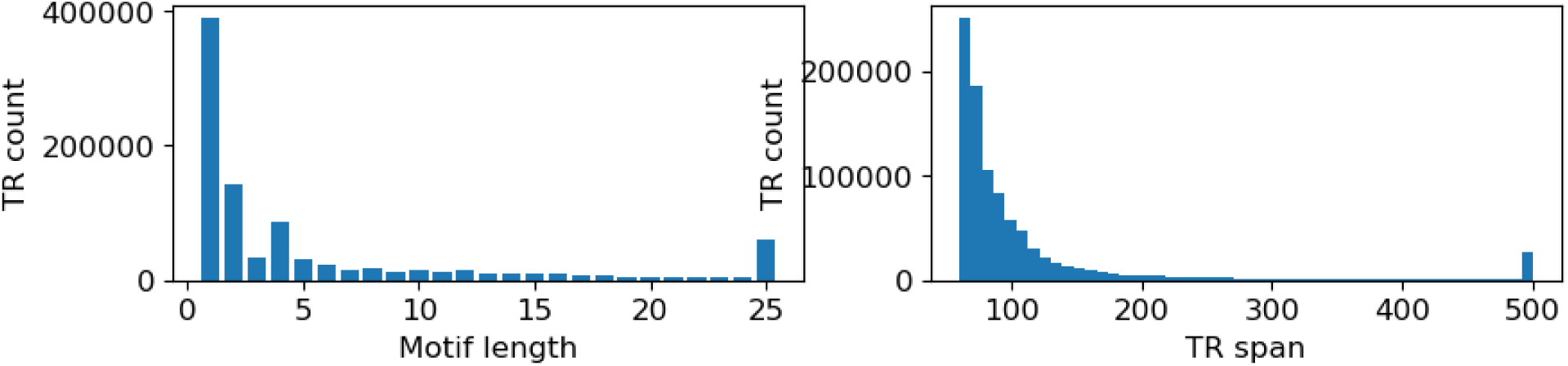
(Left) Number of TRs stratified by motif length. (Right) Distribution of TR reference spans.

**Figure S2:**
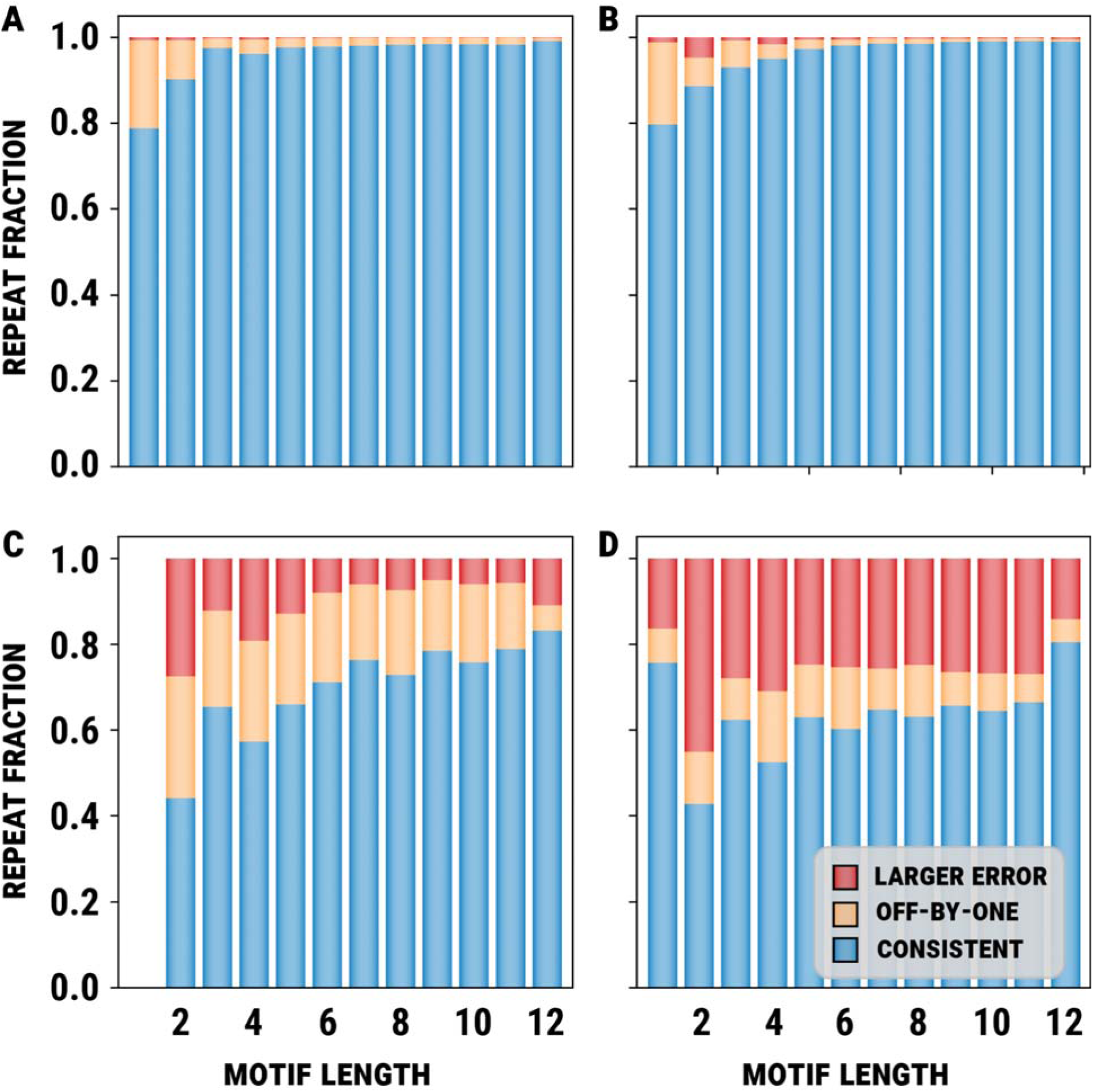
Mendelian errors in the family trio consisting of HG002, HG003, and HG004 samples for (A) TRGT, (B) tandem-genotypes, (C) Straglr, and (D) GangSTR.

**Figure S3:**
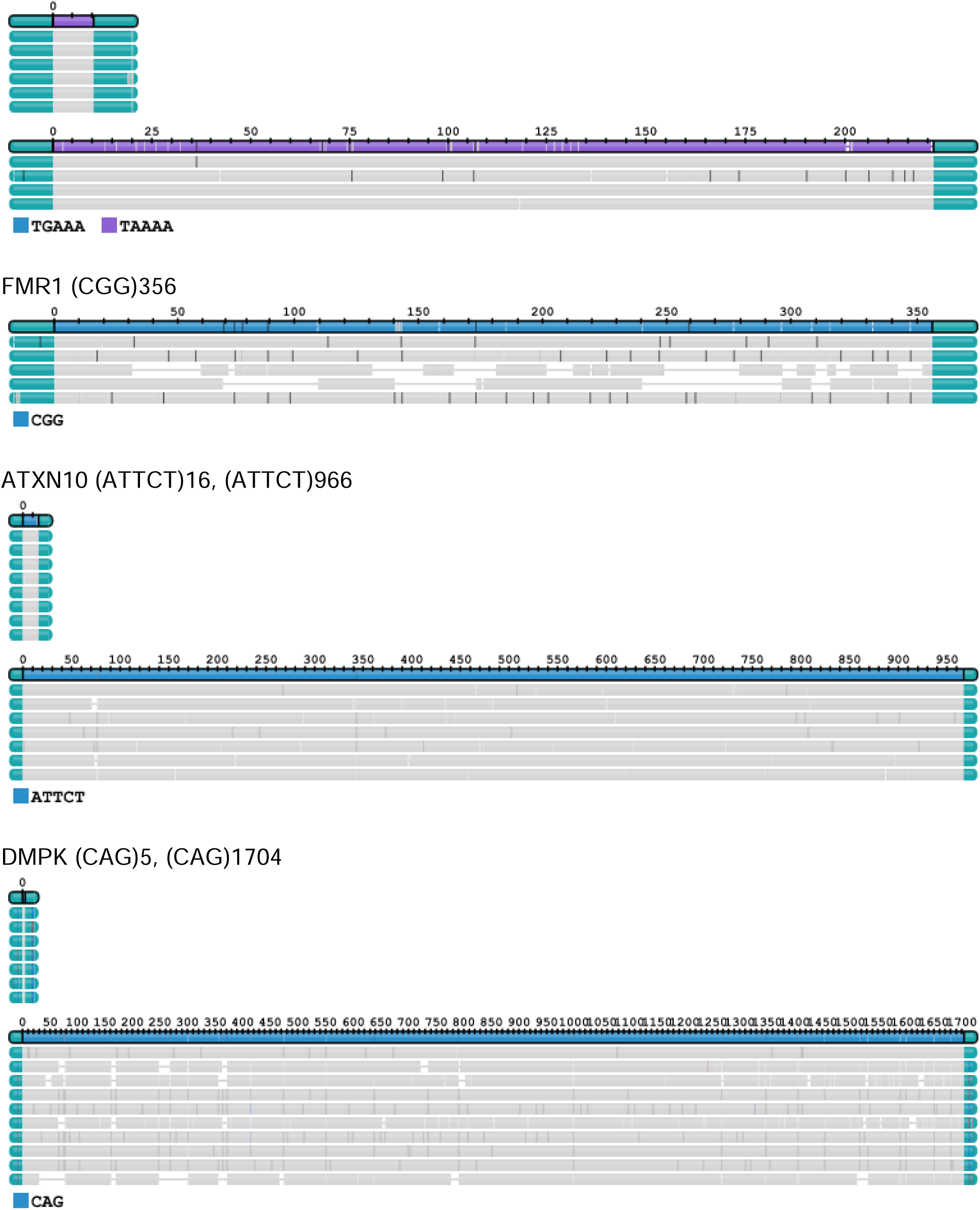
TRVZ output for select known pathogenic expansions.

**Figure S4:**
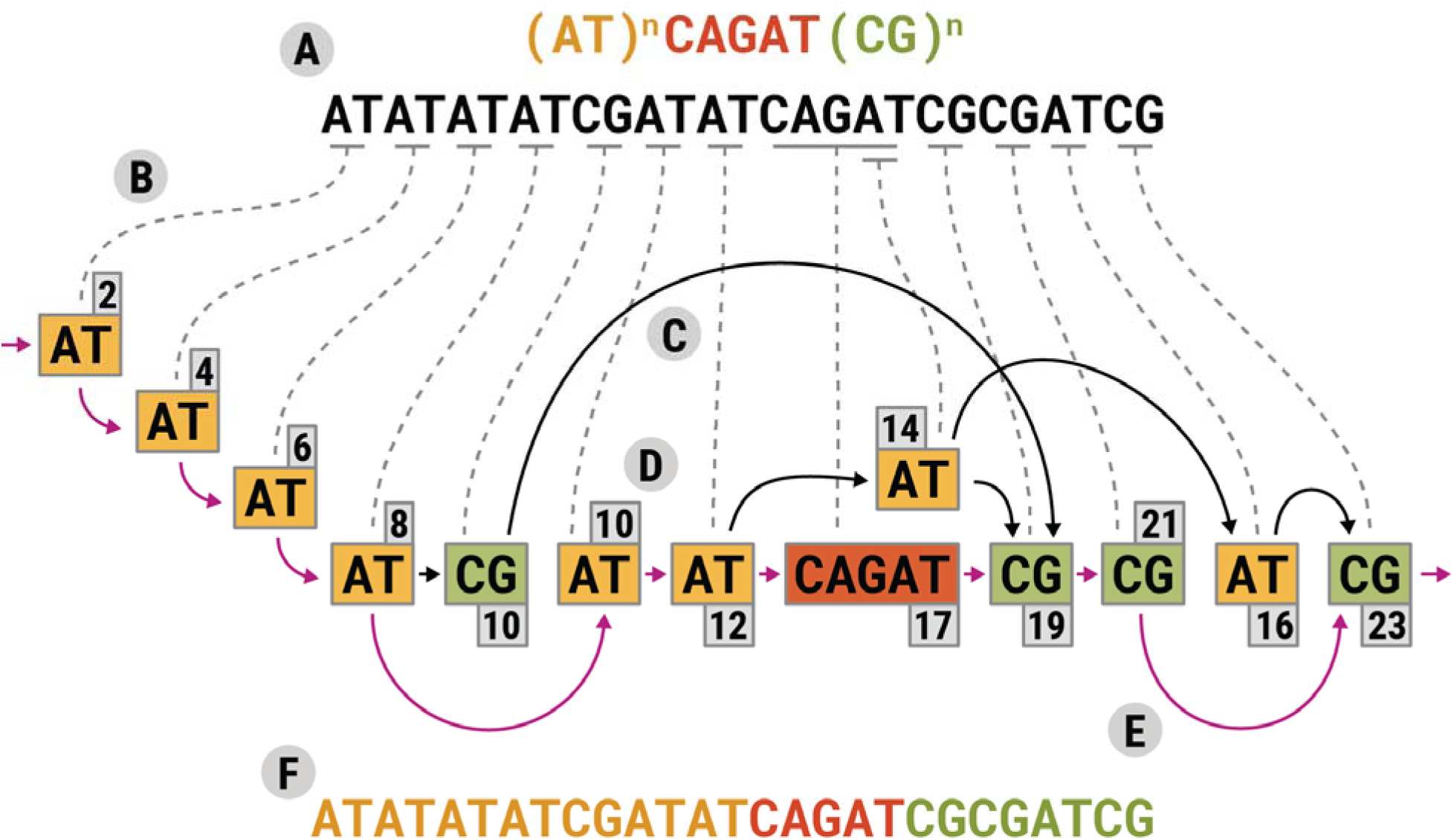
Illustration of the segmentation algorithm for simple repeats. (A) A locus definition and a query sequence. (B) Correspondence between the query sequence and graph nodes. (C) Graph edges connect nodes that are compatible with the locus definition. (D) A node score. (E) An edge of the top-scoring path. (F) Segmentation corresponding to the top-scoring path.

**Figure S5:**
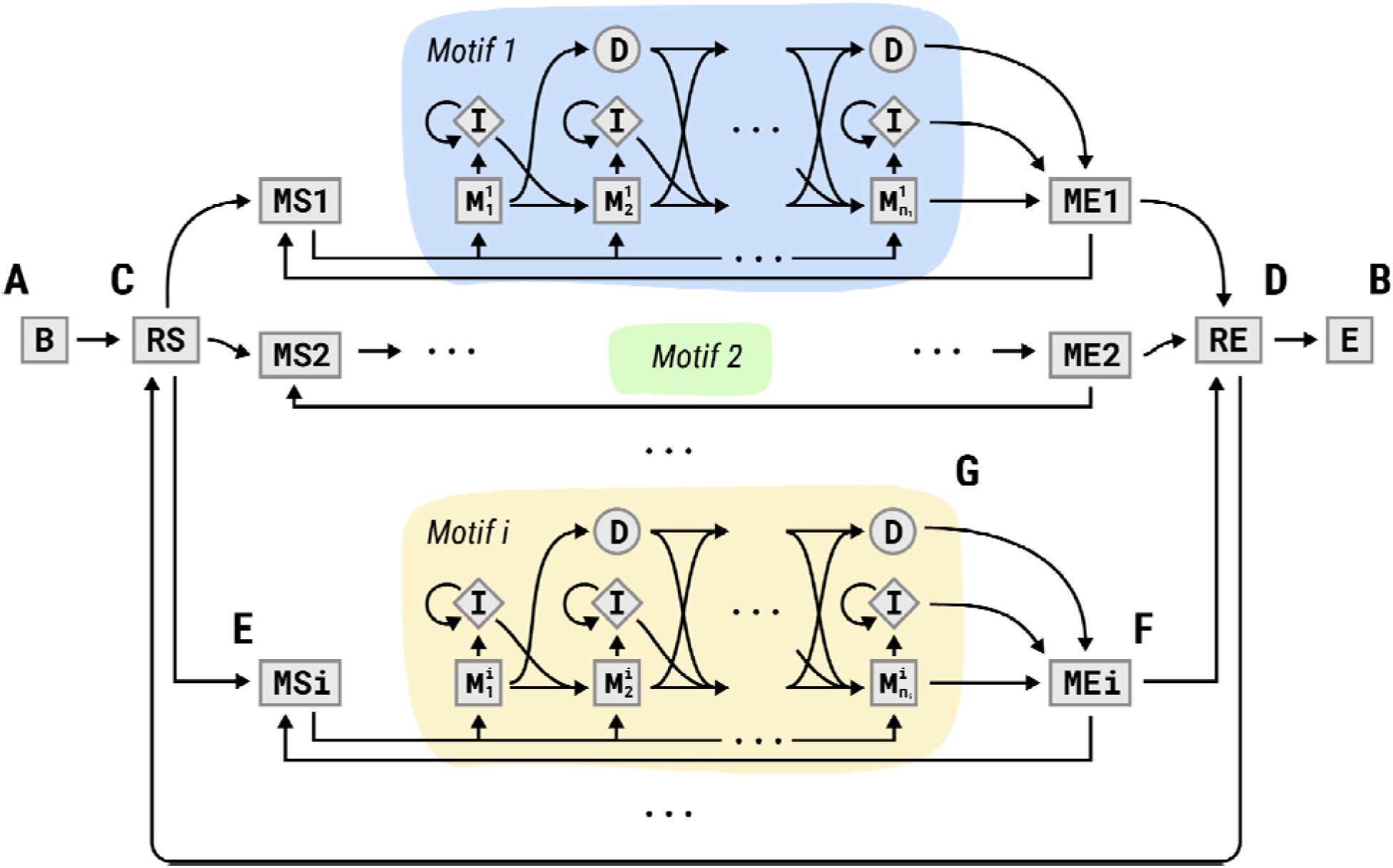
Topology of hidden Markov models corresponding to sequences consisting of runs of multiple motifs. (A, B) Start and end states. (C, D) Motif run start and end states. (E, F) Motif occurrence start and end states. (G) States corresponding to the motif occurrence sequence.

**Figure S6:**
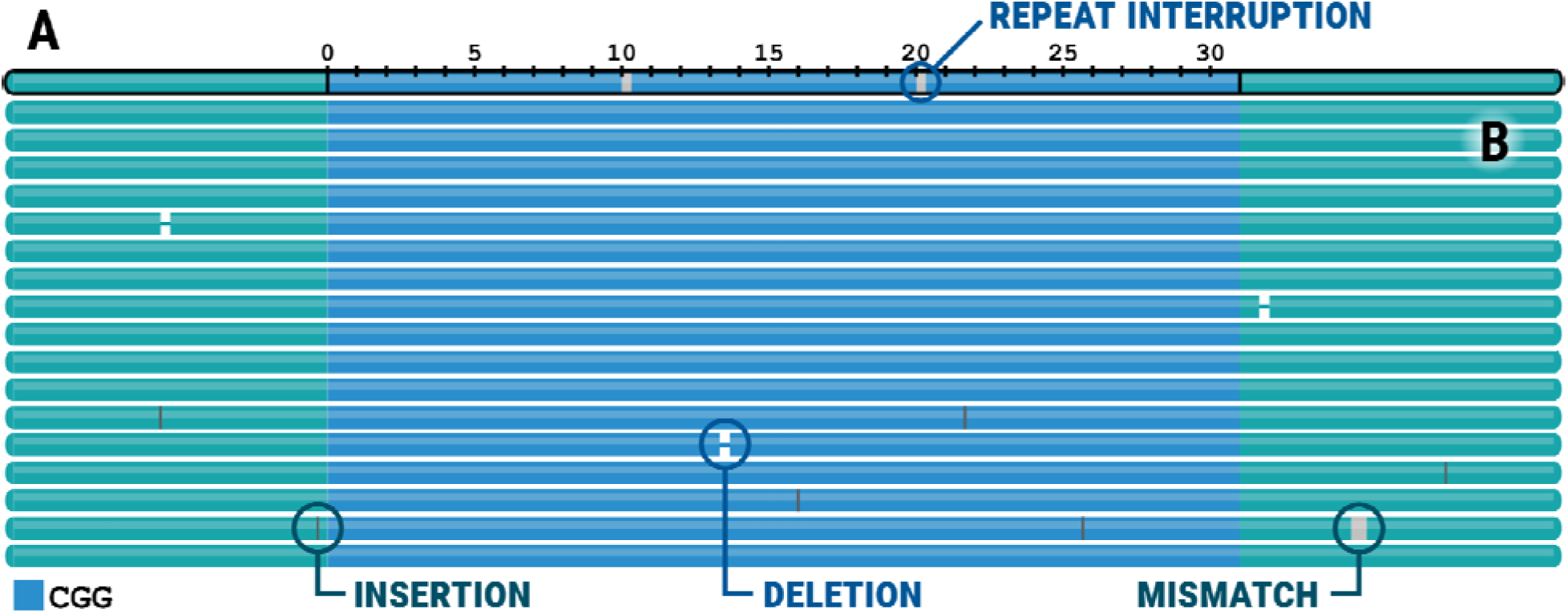
A TRVZ plot depicting the *FMR1* repeat in HG002 sample. (A) The top track depicts the consensus allele sequence (blue) and surrounding flanking sequence (green). (B) The tracks below the top track depict alignments of reads to the consensus.

**Table S1:** The list 104 HiFi samples from Genome in a Bottle (GIAB) and the Human Pangenome Reference Consortium (HPRC).

|  |  |  |  |  |  |  |  |
| --- | --- | --- | --- | --- | --- | --- | --- |
| HG001 | HG00706 | HG01255 | HG01978 | HG02145 | HG02630 | HG03098 | HG03816 |
| HG002 | HG00733 | HG01346 | HG01981 | HG02148 | HG02647 | HG03453 | HG03825 |
| HG003 | HG00735 | HG01358 | HG01993 | HG02257 | HG02656 | HG03471 | HG03831 |
| HG00423 | HG00738 | HG01433 | HG02004 | HG02280 | HG02668 | HG03486 | HG03927 |
| HG00438 | HG00741 | HG01442 | HG02027 | HG02293 | HG02683 | HG03492 | HG03942 |
| HG004 | HG007 | HG01496 | HG02055 | HG02300 | HG02698 | HG03516 | HG04115 |
| HG00544 | HG01071 | HG01884 | HG02071 | HG02486 | HG02717 | HG03540 | HG04157 |
| HG005 | HG01099 | HG01887 | HG02074 | HG02523 | HG02723 | HG03579 | HG04160 |
| HG00609 | HG01106 | HG01891 | HG02080 | HG02559 | HG02738 | HG03654 | HG04184 |
| HG00621 | HG01109 | HG01928 | HG02083 | HG02572 | HG02809 | HG03669 | HG04187 |
| HG00642 | HG01123 | HG01934 | HG02109 | HG02602 | HG02818 | HG03688 | HG04199 |
| HG00673 | HG01175 | HG01943 | HG02132 | HG02615 | HG02886 | HG03710 | HG04204 |
| HG006 | HG01243 | HG01952 | HG02135 | HG02622 | HG02970 | HG03804 | HG04228 |

**Table S2:** Individuals with known genotypes at pathogenic STR loci. Clinical testing was performed by triplet-primed PCR (TP-PCR), traditional PCR and/or Southern blot.

| STR locus | Chr | Pathogenic expansion | Sex | Clinical test | Clinically reported result | TRGT call | Notes |
| --- | --- | --- | --- | --- | --- | --- | --- |
| DMPK | chr19 | (CAG)50+ | XX | Southern Blot | ~1600 and 5 repeats | (CAG)5,<br>(CAG)1704 | Shows expected methylation |
| STARD7 | chr2 | (TGAAA)>340<br>(TAAAA)>274 | XY | TP-PCR | Expanded >200 and normal (10-30 repeats) | (TAAAA)10,<br>(TAAAA)221 |  |
| STARD7 | chr2 | (TGAAA)>340<br>(TAAAA)>274 | XX | TP-PCR | Expanded >200 and normal (10-30 repeats) | (TAACA)13,<br>(TAAAA)224 | Alternate motif in shorter allele |
| FMR1 | chrX | (CGG)>200 | XY | TP-PCR | >200 repeats | (CGG)356 |  |
| FMR1 | chrX | (CGG)>200 | XX | TP-PCR | Two alleles <200, not positive for full mutation | (CGG)29,<br>(CGG)117 | Premutation |
| ATXN10 | chr22 | (ATTCT)800+ | XY | PCR,<br>Southern Blot | 1071 and 16 repeats | (ATTCT)16,<br>(ATTCT)966 |  |

